# Evidence that recent climatic changes have expanded the potential geographical range of the Mediterranean fruit fly

**DOI:** 10.1101/2023.11.15.566983

**Authors:** Anna M. Szyniszewska, Hanna Bieszczak, Karol Kozyra, Nikos T. Papadopoulos, Marc De Meyer, Jakub Nowosad, Noboru Ota, Darren J. Kriticos

## Abstract

The species distributions migration poleward and into higher altitudes in a warming climate is especially concerning for economically important insect pest species, as their introduction can potentially occur in places previously considered unsuitable for year-round survival. We explore the expansion of the climatically suitable areas for a horticultural pest, the Mediterranean fruit fly (medfly) *Ceratitis capitata* (Diptera, Tephritidae), with an emphasis on Europe and California. We reviewed and refined a published CLIMEX model for *C. capitata*, taking into consideration new records in marginal locations, with a particular focus on Europe. To assess the model fit and to aid in interpreting the meaning of the new European distribution records, we used a time series climate dataset to explore the temporal patterns of climate suitability for *C. capitata* from 1970 to 2019. At selected bellwether sites in Europe, we found statistically significant trends in increasing climate suitability, as well as a substantial northward expansion in the modelled potential range. In California, we also found a significant trend of northward and altitudinal expansion of areas suitable for *C. capitata* establishment. These results provide further evidence of climate change impacts on species distributions and the need for innovative responses to increased invasion threats.

## Introduction

Climate is the principal factor defining the potential distribution of poikilotherms ^1^. Hence, climatic changes are expected to impact the geographic ranges of a variety of organisms, including pests affecting their ability to overwinter in new areas, exposure to heat or cold stresses, and the number of generations they are able to complete within a season ^2^. As these organisms spread beyond their native ranges, they may endanger natural and agricultural ecosystems, often triggering expensive management responses.

Climate change manifests through substantial shifts in seasonal, interannual and inter-decadal weather patterns. Hence, the identification of linkages between climate change and biological trends in this context is difficult. This challenge is further compounded for invasive species because they take time to expand their range to occupy their climatic niche. These changes may be short in duration and further interrupted by the return of less-favourable conditions. In the long term, favourable conditions may prevail, leading to successful range expansion. Bioclimatic niche models can be useful in revealing these trends and attributing them to climate change ^3^.

The Mediterranean fruit fly *Ceratitis capitata* (Wiedemann) (Diptera: Tephritidae) is a notorious invasive species reported to be among the world’s most economically important pests ^4^. It is a highly polyphagous insect, feeding on over 300 plant species, including major commodities such as citrus, peach, persimmon, apple and mango ^5^. *Ceratitis capitata* has a wide geographical distribution and is well adapted to a broad range of climates ^6,7^. It has spread from its native origin in sub-Saharan Africa throughout much of the continent. Over time, it has colonised habitats far from its original range, including areas experiencing a temperate climate ^8–10^. *Ceratitis capitata* is now established in North Africa, Europe, South and Central America, Australia, and Hawaii ^11^. In the USA it is regularly detected in California, though its invasion status is hotly debated ^12,13^.

Given *C. capitata*’s economic importance, there have been many attempts to map the pest risk it poses (Table 1). The mechanistic models appear to perform better in relation to the modelled potential distribution. The correlative models mostly appear to suffer from the inclusion of ephemeral records in the training datasets.

**Table 1.** Summary of the published models of the potential distribution of *Ceratitis capitata*.

| Model type | Source | Model and met data | Characteristics | Evaluation of results |
| --- | --- | --- | --- | --- |
| Mech-anistic | Vera *et al.*<sup>14</sup> | CLIMEX<br>Met data: CLIMEX in-built met station data and gridded Climate Research Unit (CRU) data. Average climatology for 1961-1990. | Trained on distribution records in Argentina and Australia, estimating global potential distribution. Included consideration of the effects of irrigation. | Model indicated a maximum poleward establishment limit of Genova (north-western Italy). May have been slightly conservative in the estimated range extent. |
|  | Gutierrez and Ponti<sup>12</sup> | CASAS<br>Max-min temps and solar radiation data from UC DAVIS and UCEA. Time range not specified (1996-2006) | Estimated the invasive potential of *C. capitata* in California and Italy. | The overall pattern of response in Italy appears plausible, though model results elsewhere make it impossible to judge the fit generally. |
|  | Gilioli *et al.*<sup>18</sup> | PBDM<br>NCDC NOAA (2020) Climate change scenarios (2030 and 2050) | Explored the distribution limits of *C. capitata* in Europe under current and future climate scenarios | Appears to do a good job of modelling the northern range limit for establishment of *C. capitata* in Europe. |
| Cor-rela-tive | De Meyer *et al.*<sup>15</sup> | GARP | Trained on both native and non-native distribution data. | Modelled most of Europe, including parts of Scandinavia as being suitable for establishment. Appears to substantially overestimate the potential for establishment in cold environments. |
|  |  | PCA<br>WorldCLIM 1950-2000 |  |  |
|  | Li *et al.*<sup>16</sup> | MaxEnt<br>WorldCLIM 1950-2000 | Focused on China. Trained on distribution records that included ephemeral populations in northern France and also on location records from xeric locations where irrigation is practised. | Northern Scotland and parts of Greenland were modelled as suitable for *C. capitata* to establish (implausibly cold), and large swathes of the Sahara Desert and most of Central Australia (implausibly dry). Also produced some Moiré modelling artefacts in the suitability patterns that are unrelated to topoclimatic variables. |
|  | Szyniszewska and Tatem<sup>17</sup> | MaxEnt<br>WorldCLIM: 1950-2000<br>20-year time series of AVHRR Land Surface Temperatures | estimate seasonally varying suitability for *C. capitata* based on a comprehensive set of spatio-temporal occurrence data and bioclimatic variables for 1950-2000 | One or two outlying occurrence records at the base of the Italian Alps have probably anchored the response functions, leading to the implausible projection that north-western Europe as far north as Denmark is suitable for persistence of *C. capitata*. Undoubtedly, some of these training records were from transient populations. |

To improve our understanding of the climate suitability patterns of *C. capitata* globally and to explore recent reports of its dynamics in southern Europe, we revised the Vera *et al*.^14^ CLIMEX model, refitting parameter values according to the available literature and a broader range of occurrence points and seasonal *C. capitata* trapping data excluding points that could represent transient populations from model fitting. We run composite irrigation simulations including top-up irrigation in irrigated zones, and natural rainfall conditions elsewhere, to illustrate areas suitable for *C. capitata* establishment and ephemeral occupation. We examine the trends in interannual climate suitability index values returned by CLIMEX to identify areas where the trends are statistically significant.

## Results

### Ecoclimatic suitability

The potential global distribution of *C. capitata* spans tropical, subtropical and warm temperate zones (Figure 1). The main limiting factors in their distributions are the cold stresses observed in higher temperate latitudes and the dry stress in non-irrigated areas of subtropical climates. The potential global distribution when 2.5 mm irrigation is added as a top-up to natural rainfall extends *C. capitata* potential distribution notably in drier climates of North America, North Africa, the Middle East and South Asia. Figure 2 produced with QGIS software illustrates a composite scenario: model outputs with a top-up irrigation in areas classified as equipped with irrigation and rainfed in areas classified as not equipped with irrigation^15^.

**Figure 1.**
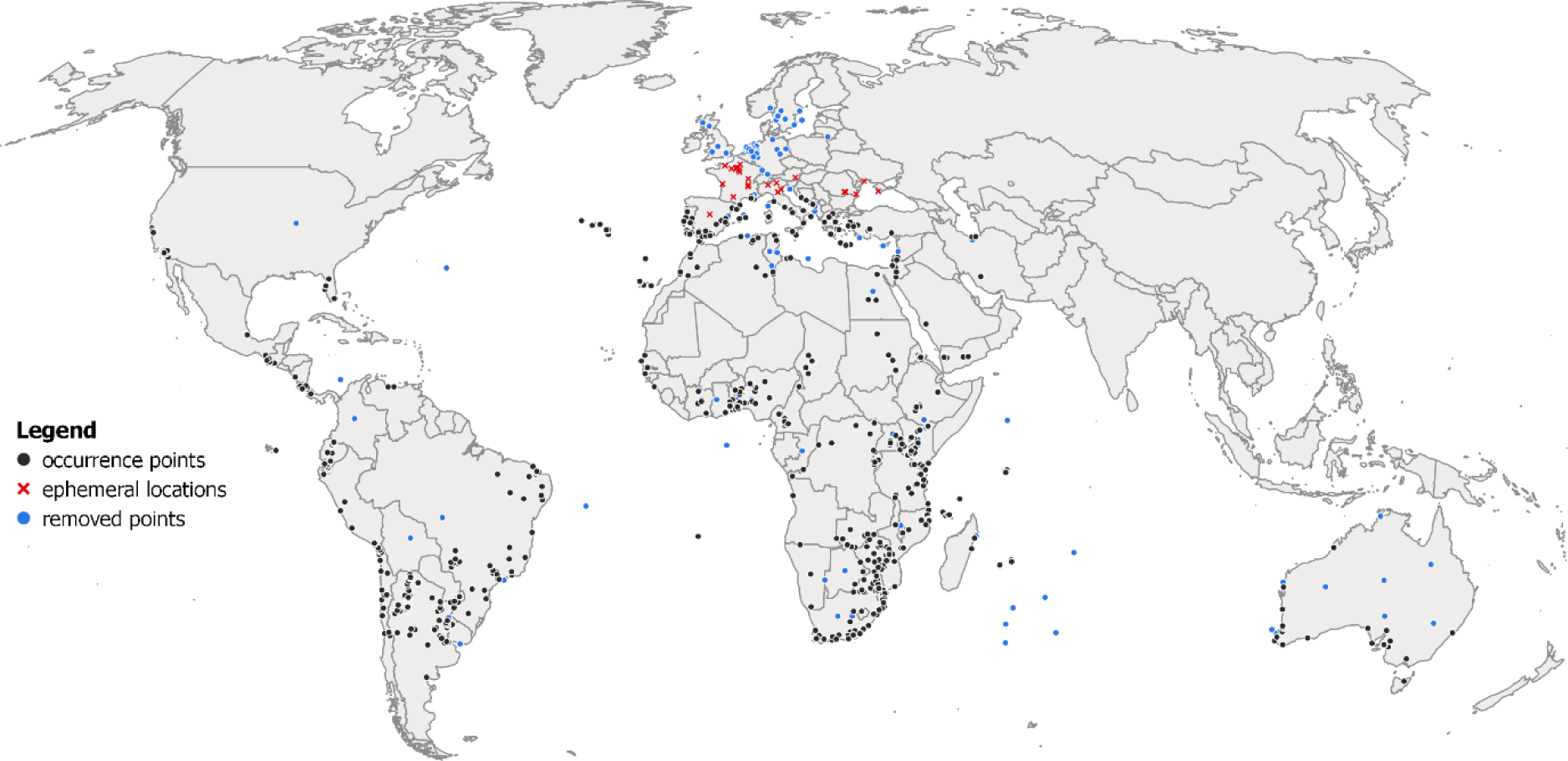
*Ceratitis capitata* historic occurrence data that was used for fitting the CLIMEX model. While most records represent locations where *C. capitata* is established, some represent historical records where the ability of fly populations to overwinter was equivocal (red crosses and blue dots). Points that did not pass inclusion criteria in our cleaning stage or were located in areas where there is no evidence of medfly ability to establish were removed from the analysis (blue dots). The map lines do not necessarily depict accepted national boundaries. The figure was created with QGIS version 3.22.9 (www.qgis.org).

**Figure 2.**
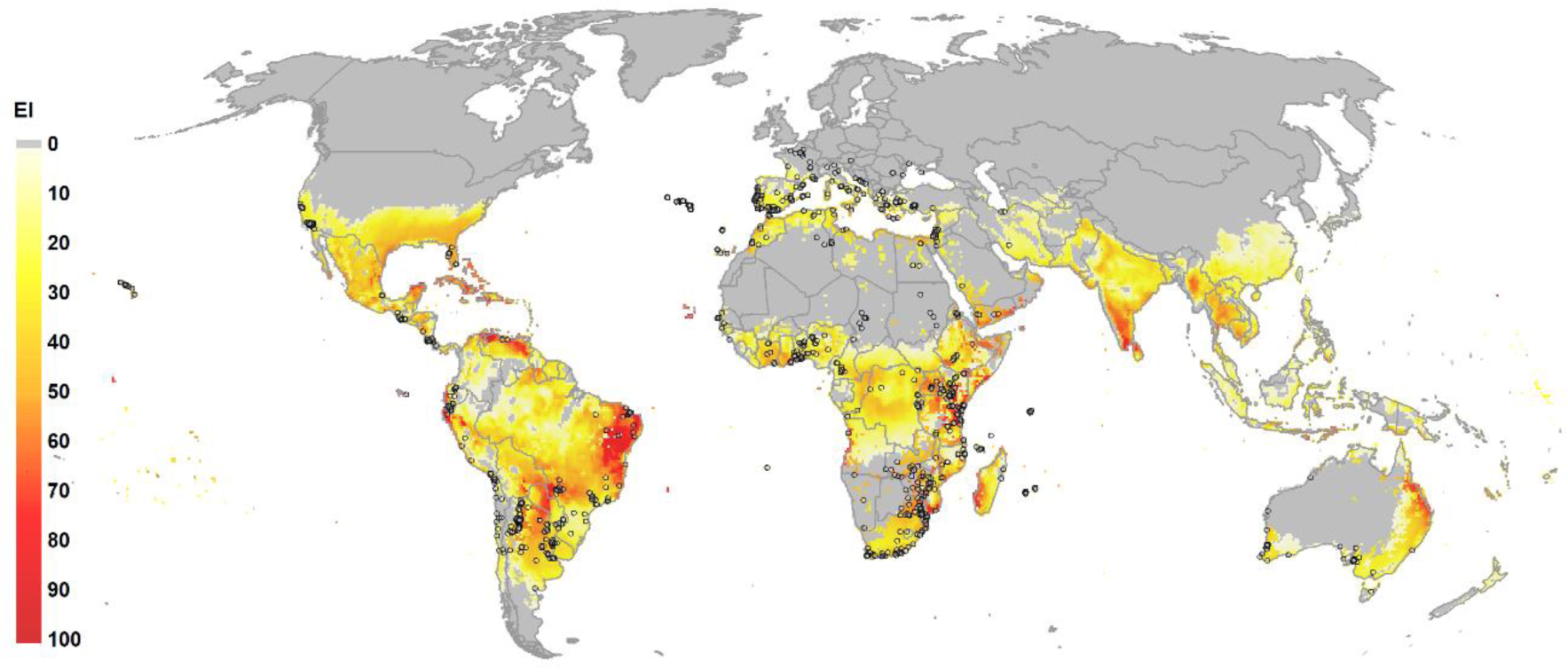
The Compare Locations model in CLIMEX 4.1 run with 30-year average climate data centred on 1995 (CM30 1995H) to produce global Ecoclimatic Index (EI) map for a composite rainfed and irrigated scenario based on a set of parameters presented in Table 2. Circles indicate the collated occurrence records of medfly globally. The map lines do not necessarily depict accepted national boundaries. The figure was created with QGIS version 3.22.9 (www.qgis.org).

**Table 2.**
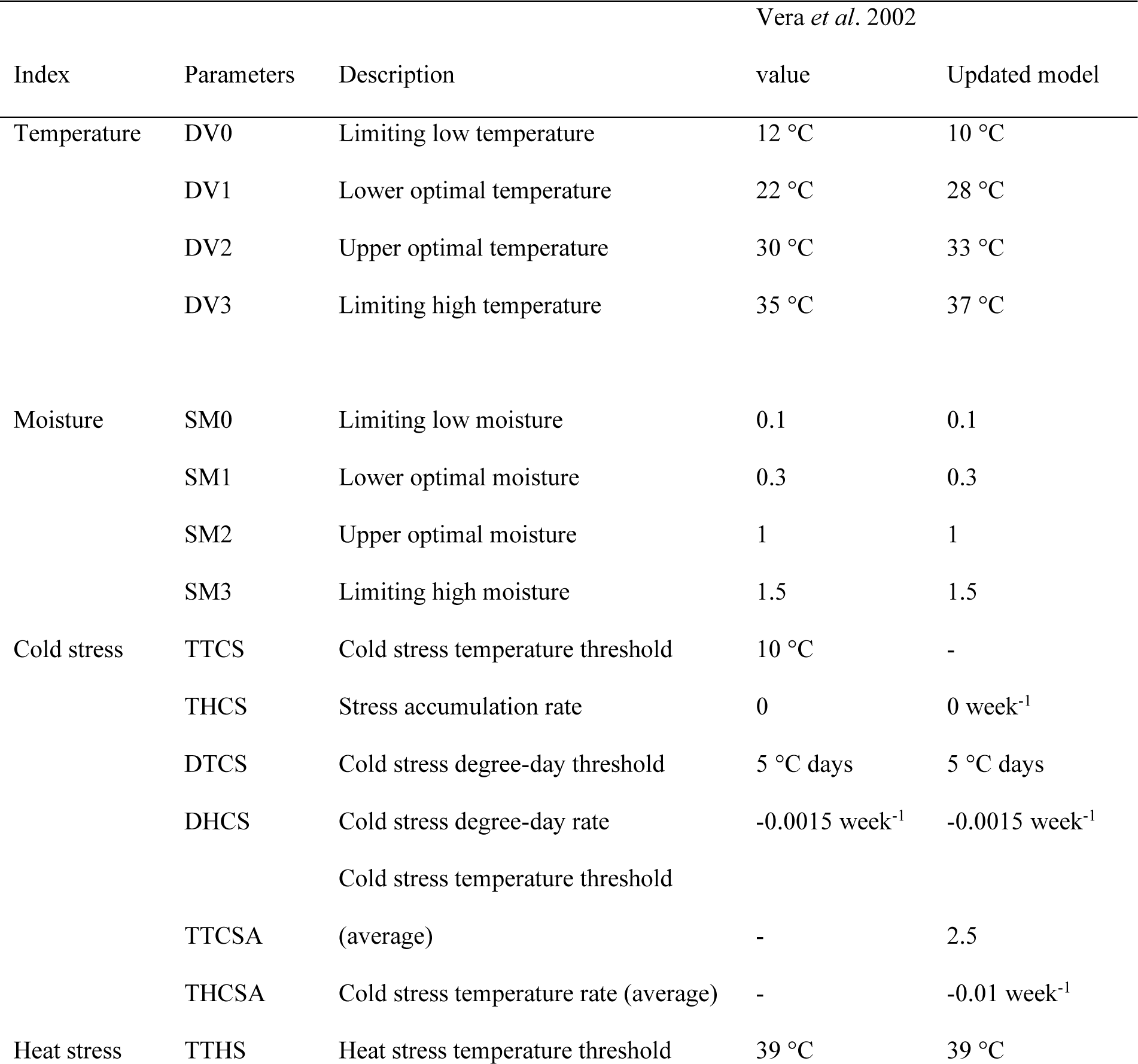
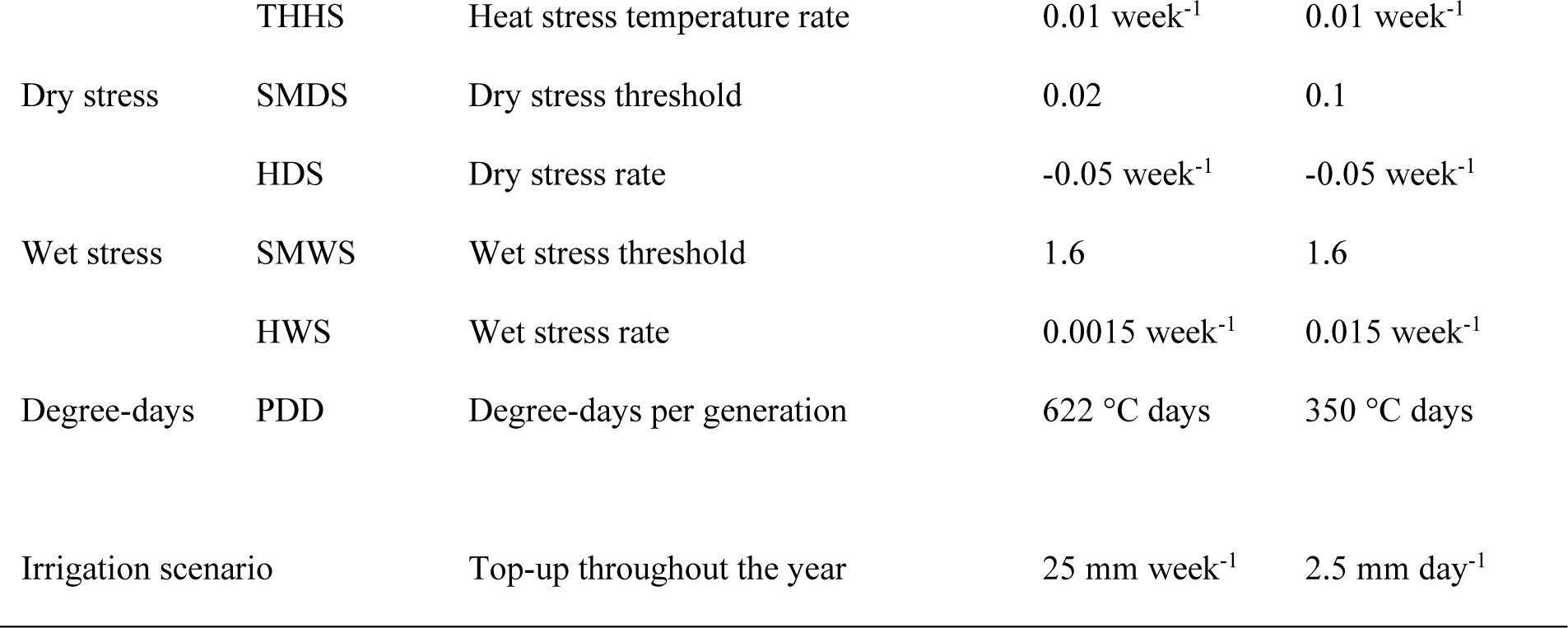
CLIMEX parameters used to model the distribution of *Ceratitis capitata*. Parameters for a previously published CLIMEX model. ^14^ **are presented for comparison.**

### CLIMEX index values in 1970-1979 and 2010-2019

Comparison of the average CLIMEX index values in 1970-1979 against 2010-2019 revealed increase in the overall climatic suitability (Ecoclimatic Index – EI) in many areas of the globe including Europe and California (Figures 3, S3, S4). Overall annual Growth Index values (GI) increased significantly in Europe, notably due to increases in Temperature Index (TI). Cold Stress (CS) values below 100 which is a threshold is observed across larger southern areas of Europe in the recent decade. No notable changes are observed for the Moisture Index (MI) and Heat Stress (HS) values.

**Figure 3.**
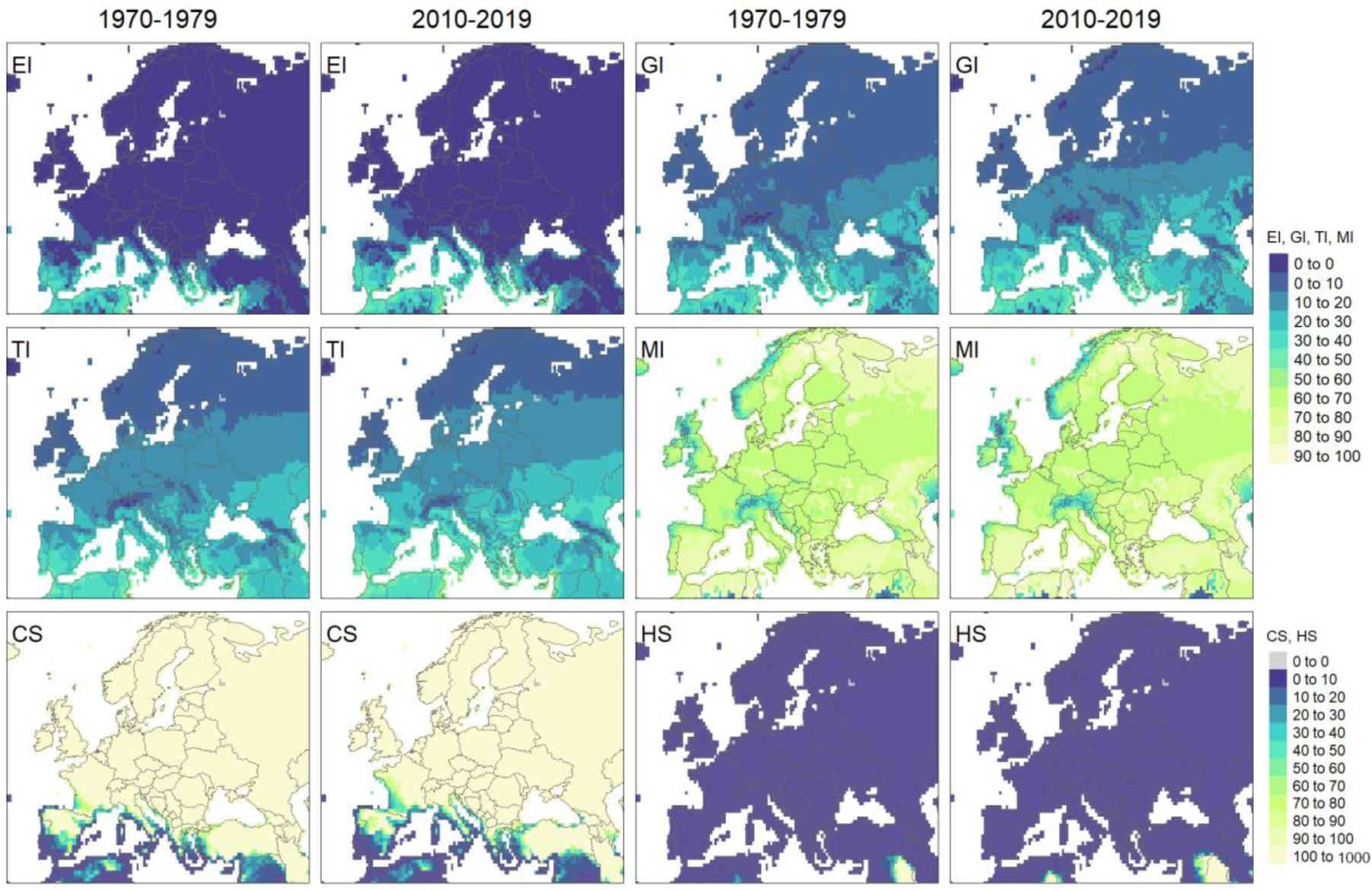
CLIMEX model results representing 10-year average values of Ecoclimatic index (EI), Growth Index (GI), Temperature Index (TI), Moisture Index (MI), Cold Stress (CS) and Heat Stress (HS) in Europe for 1970-1979 (1^st^ and 3^rd^ column) and 2010-2019 (2^nd^ and 4^th^ column). The Compare Locations/Years function was run utilising CRU climate data returning annual index and stress values. Figure was produced with tmap^16^ packages in R^17^.

### Rate of change in climatic suitability

The linear trend fitted to every grid cell of the timeseries of the model runs for 1970-2019 presents the annual rate of change in CLIMEX indices values (Figures 3, S5, S6) in areas where the trend is significant (*p <* 0.05; Figures 4, S7, S8, S9). Notably, we observe increasing EI values in vast areas of Mediterranean Europe. This increase appears driven by increasing Growth Index at the rate of up to slightly above 0.5 in many areas of southern and central Europe. TI has a positive rate of change of over 0.1 in all areas of Europe except northern and eastern parts. CS is decreasing at a highest rate in central, southern and some parts of northern Europe. The MI changes are positive in most of southern Europe and negative in the north. Heat stress values changes are negligible on the European continent. In large areas of southern Europe, the trend in EI changes increase was significant (Figures 4, S7).

**Figure 4.**
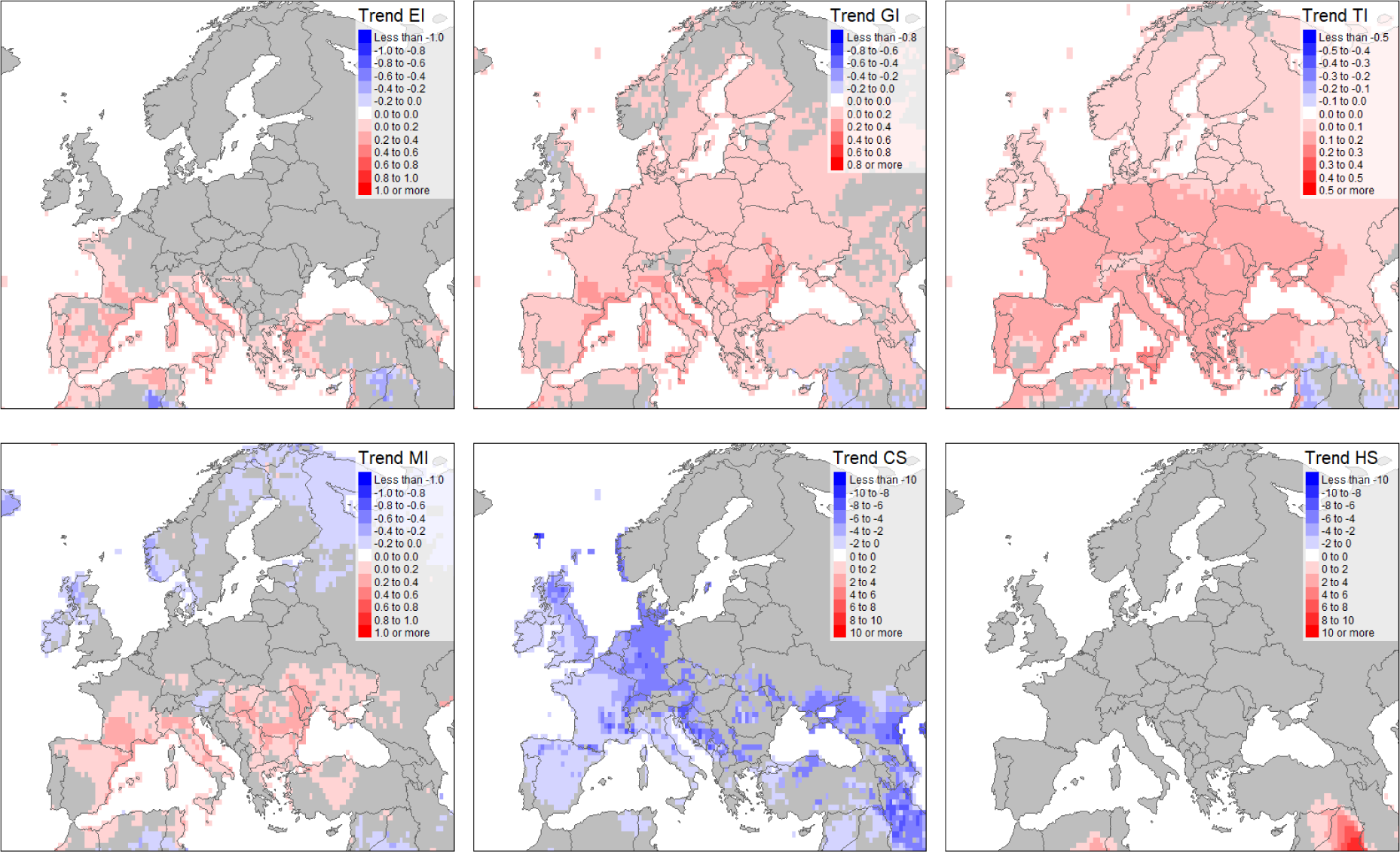
Annual rate of change (trend) values returned by lm() function in R statistical programming software run on annual values returned by Compare Locations/Years function for 1970-2019 for six CLIMEX indices: Ecoclimatic index (EI), Growth Index (GI), Temperature Index (TI), Moisture Index (MI), Cold Stress (CS) and Heat Stress (HS). Figure was produced with tmap^16^ packages in R^17^.

Positive TI changes were significant across the majority of Europe. GI changes in the north, as well as small patches of southern Europe, including areas in Spain and Austria, had a significant trend of positive changes. Changes in the MI appear more complex, with more significance in either the south or northern parts of the continent. CS trends in western Europe appear significant, in contrast to eastern Europe, where this trend’s p values were well above 0.05.

We chose eight bellwether locations in Europe, and three locations in California to investigate the trends in climatic suitability (EI, GI, TI, MI) and stresses limiting species ability to survive (CS and HS) (Figure 5). Chosen locations in Spain experience an increasing significant positive trend in the suitability for *C. capitata* (Valencia = 0.35 and Barcelona = 0.34). In France, Marseille has increased its overall EI from nearly 0 to around 15 in recent years, with the trend of over 0.3 annual increase in EI since 1970s. Constantia, in Romania, experienced only a few years during which *C. capitata* would have been capable of overwintering (EI > 0.5). Locations in Po Valley, Split and Thessaloniki continue to experience increasingly favourable conditions enabling *C. capitata* to overwinter and thrive in these areas. In California, we observed statistically significant changes in EI only in San Diego, Los Angeles and San Francisco (Figure 5). The rate of annual change in EI in these locations is 0.1 in San Diego, and 0.7 in Los Angeles and San Francisco.

**Figure 5.**
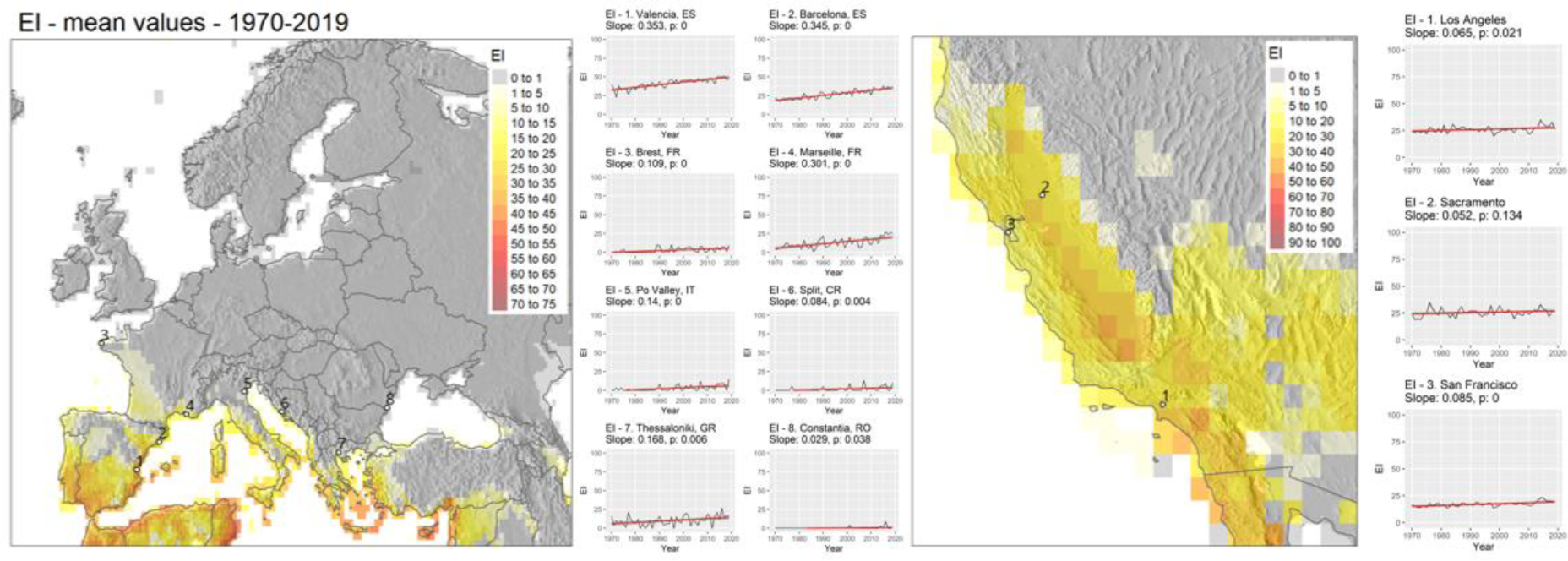
Representation of 50 years (1970-2019) average EI values in Europe and California calculated using annual timeseries of Ecoclimatic Index (EI) values returned by Compare Locations/Years function in CLIMEX using CRU climate data 50-years time-series. Annual EI values for selected locations in Europe and California have a linear trend fitted. Slope values represent the annual trend of change in index values over model outputs timeseries represented by a fitted linear model, and *p value* represents associated significance value of the slope parameter. Figure was produced with tmap^16^ and ggplot^18^ packages in R^17^.

Globally, we observe poleward expansion in the areas suitable for medfly (EI 0.5 or above) in the temperate latitudes (Figure 6). This trend, illustrating the difference in average irrigation composite 10-year EI between 1970-79 and 2010–19, is driven mostly by reduced CS and, consequently, an increasing capacity for the organism to overwinter in these locations. Conversely, in subtropical regions, certain areas that were once suitable for the medfly transitioned to unsuitable conditions. This change can be attributed to the escalation of HS and the prolonged periods of DS in regions not equipped with irrigation. In Europe, in the course of last 50 years, large areas of northern Spain, France, northern Italy, and further north into the Balkans were observed according to the outputs of our model. In California, and more broadly in the USA, the composite small area near Baja California coast appears no longer suitable, whilst the suitability appears improved across nearly the entire northern limit of species climatic niche limits.

**Figure 6.**
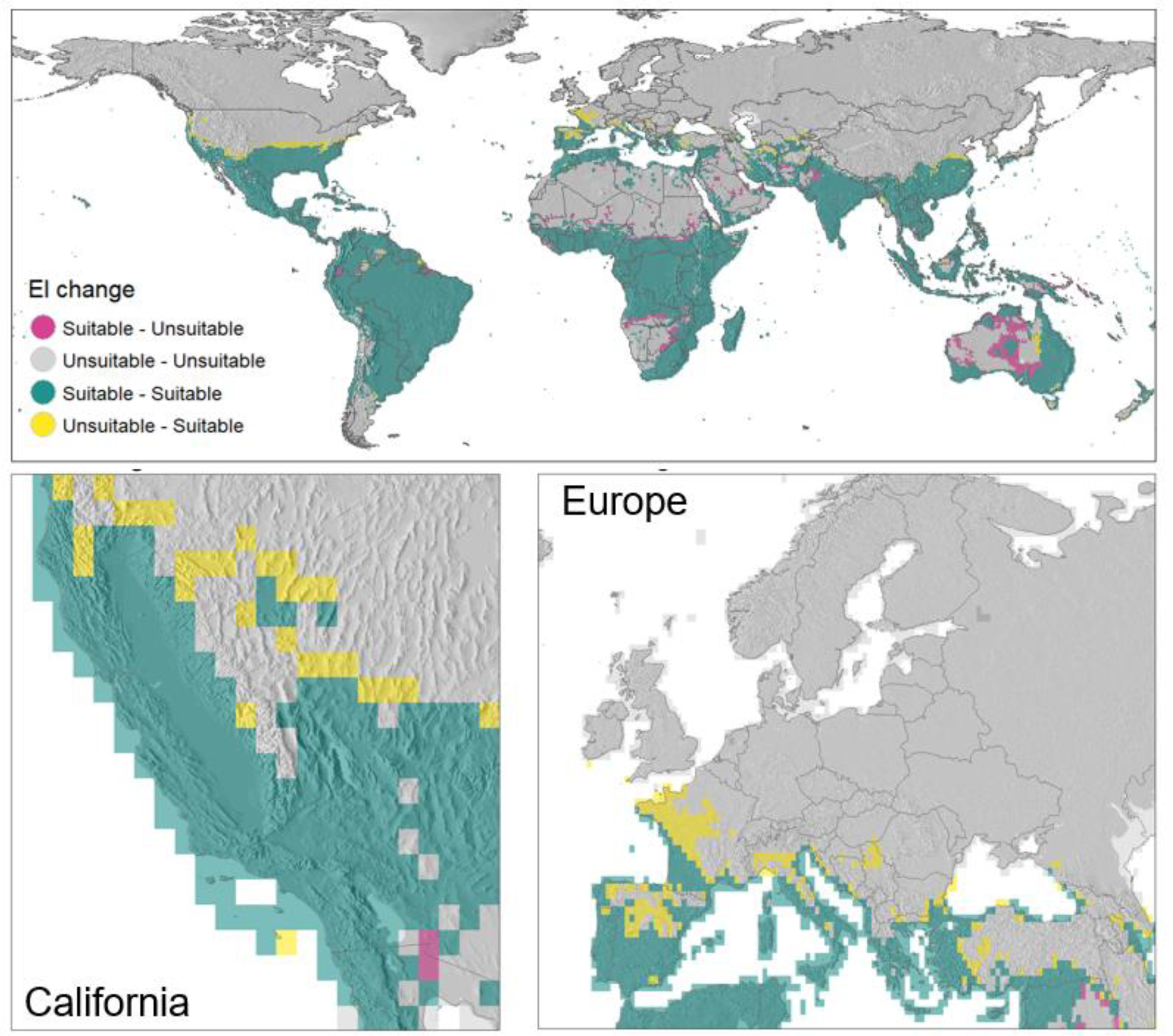
Cross-comparison of average climatic suitability values for Ceratitis capitata for 1970-1979 and 2010-2019. 10- year average EI values for each decade were calculated. Where 10-year average EI was below 0.5, the area was classified as unsuitable. Areas with EI above 0.5 were classified as suitable. Yellow highlights areas unsuitable for *C. capitata* in the 1970’s but most recently becoming suitable. Pink areas were previously suitable but are become unsuitable in recent years. Green highlights areas previously and most recently classified as suitable. Figure was produced with tmap^16^ packages in R^17^.

The examination of the changes in EI values over a 50-year period, where significant trends are observed, provides valuable insights into the impact of these changes based on latitude (Figure 7). By analysing the cumulative changes in EI across different latitudes during that timeframe, a clear pattern emerges, indicating substantial increases in EI, particularly in the temperate regions of the northern hemisphere which possess a larger landmass compared to the southern hemisphere. Although the total positive changes in EI are smaller in the higher latitudes of the southern hemisphere (over 30 degrees South), they still dominate. On the other hand, a higher proportion of negative sum of changes in EI are observed in latitudes between the equator and the tropics of the northern hemisphere, where the Sahel region and arid areas of North America, South Asia and the Middle East contribute to this trend. In the southern hemisphere, the Mato Grosso region in Brazil, as well as areas in Australia, exhibit negative trend.

**Figure 7.**
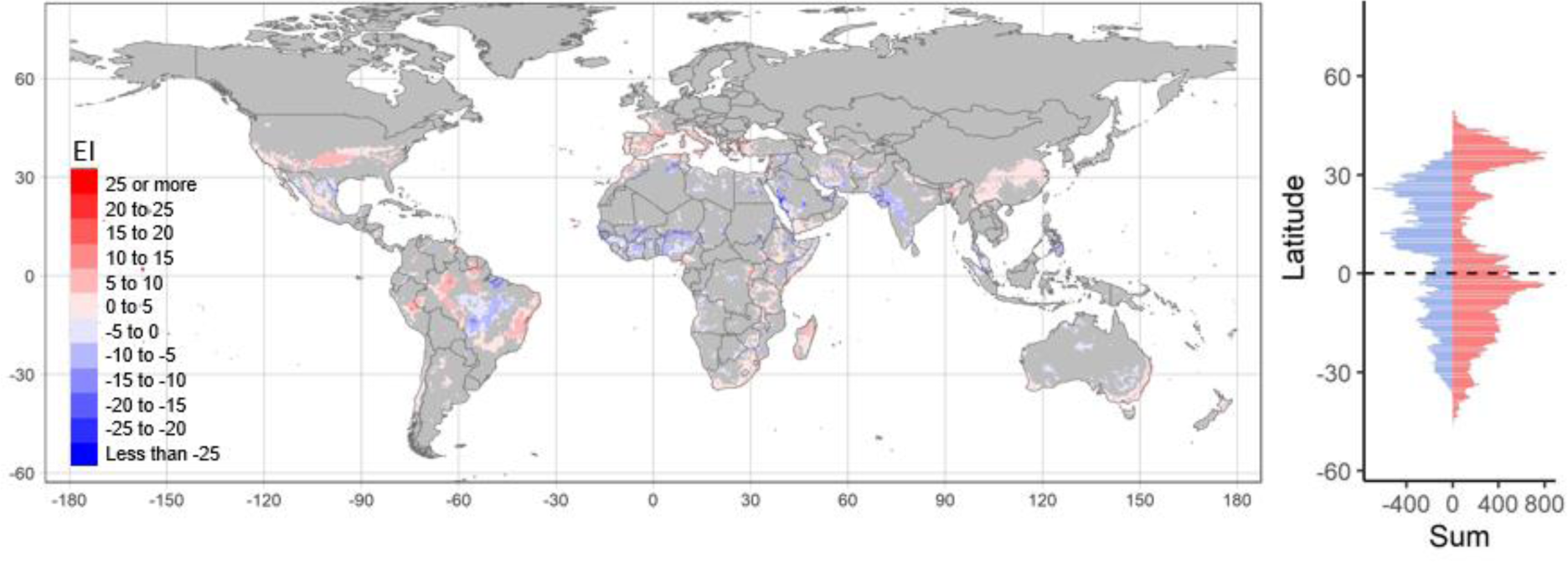
Global 50-year significant changes in Ecoclimatic Index values for *Ceratitis capitata* increase (red) or decrease (blue) between 1970 and 2019. The significant values (*p* < 0.5) are summarised across the latitudes on the right-hand panel. Figure was produced with tmap^16^ and ggplot^18^ packages in R^17^.

### Parameter sensitivity and model uncertainty

The parameter sensitivity analysis revealed little cause for concern (Table S1). There are only four parameters that had a sensitivity of greater than 1% in relation to the Range Change variable. The most sensitive variable was the dry stress threshold (SMDS) at 15.75% and the closely-related limiting low moisture parameter (SM0, 1.27%). These parameters are anchored by the permanent wilting point, which is highly stable across a wide range of host plants, and hence there is little uncertainty in setting them. The dry stress accumulation rate (HDS) is fitted to distribution data and has a sensitivity of only 2.94% and the results accord with the location records from xeric locations that are not irrigated. It is possible that *C. capitata* is persisting in drier locations that have not been documented, but the extent of this is likely very small. The next most sensitive parameter for Range Change is the Cold Stress Degree-day Threshold (DTCS) at 1.05%. This parameter is fitted to distribution data in conjunction with the Cold Stress Degree-day Accumulation Rate (DHCS), which has a sensitivity of 0.75%. The potential cold stress limits are reasonably well known, especially from European data. In Australia and South Africa, despite the latitudinal limits to the landmasses, there are elevational gradients extending into areas that are still too cold for *C. capitata* to persist. Hence, we can be satisfied that the model cold stress parameters are of little concern. The model uncertainty results are illustrated in Figure S10.

## Methods

### Ceratitis capitata occurrence data

*Ceratitis capitata* occurrence data were obtained from the Global Biodiversity Information Facility (GBIF: www.gbif.org), Royal Museum for Central Africa, and published literature^19^ (Figure 1).

Records dated pre-1950 were excluded, assuming that older records may be more prone to digitisation errors, due to potentially less accurate data collection techniques. Moreover, older records might represent occurrences in habitats that changed over time. Thus, we are focusing on more recent data points. Records representing *C. capitata* detections only, and those that fell well outside the known climatic niche and no evidence of *C. capitata* overwintering and establishment were considered outliers and thus excluded from the analysis ^20^. Data were reviewed for clear outliers with incorrectly typed coordinates and those that evidently showed country or administrative units’ centroids. The identified outliers were removed from the dataset. The GBIF data contained many occurrence points from areas in northern latitudes well outside the known distribution area of *C. capitata*. We removed obvious outliers from countries where *C. capitata* is listed as absent according to EPPO and CABI Pest Distribution Compendia. For the remaining outliers located in northern locations of countries where the distribution is believed to be limited to southern areas, we consulted in-country experts to indicate which records likelyindicate true overwintering generations and which represent seasonal *C. capitata* trapping with no evidence of overwintering potential. Several bellwether sites remained, notably in Spain, France, Italy, Austria and Romania. The medfly occurrence maps were prepared using Free and Open Source QGIS software. The dataset is available in an open repository ^21^.

### Climate data

For fitting the CLIMEX model, we used the CRU TS4 data for 1970-2019 with 30’ spatial resolution^22^. The dataset contained monthly values for daily minimum and maximum temperature (°C), vapour pressure (hPA), and monthly rainfall totals (mm). For long-term averages, we used 30-years CliMond data centred on 1995 (CM10 World 1995H V2) with daily minimum and maximum temperatures, monthly rainfall totals and relative humidity (at 09:00 and 15:00 h) at a spatial resolution of 10 arc minutes.

### Irrigation

To account for irrigation, we ran the CLIMEX models with an irrigation scenario as a rainfall top-up of up to 2.5 mm day^-1^ applied continuously for 12 months. No irrigation was added if rainfall in a specific location was equal to or exceeded 2.5 mm day^-1^. We used the Global Map of Irrigation Areas (GMIA) version 5.0 to identify areas with a significant area under irrigation ^23^ (Figure S1). In the GMIA dataset the area equipped for irrigation was expressed in hectares per cell. This dataset was resampled to match the spatial resolution of the CRU TS4 dataset. For each 30’ cell, if the irrigation area is greater than 10 ha, the irrigation scenario was used for composite maps (Figure S2).

### Niche Modelling

The CLIMEX V4.1 modelling package^24,25^ (Hearne Scientific Software, www.hearne.software) was used in this study to fit an ecological niche model of *C. capitata* to the occurrence locations. CLIMEX is a hybrid inductive and deductive model estimating a species response to climate at each location based on the assumption that in the course of a year, populations may experience one or more seasons that are favourable for growth and, conversely, one or more seasons that are stressful, during which the population declines^24,25^. The model integrates weekly responses of a population to climate and calculates a set of climate suitability indices. The CLIMEX model assumes that population growth (development) occurs between the specified range of temperature values and between specified values of soil moisture parameters (SM0 - lower moisture and SM3 - higher moisture limit, see below). These growth functions are combined in accord with the Sprengel-Liebig Law of the Minimum ^26,27^ and Shelford’s Law of Toleration ^28,29^. Stresses, leading to negative population growth, are assumed to accumulate outside these development thresholds. For example, Cold Stress (CS) and Dry Stress (DS) can only begin to accumulate once the temperature or soil moisture drops below Limiting low Temperature (DV0) or Limiting Lower Moisture Limit (SM0) respectively (Table 2).

Conversely, Heat Stress (HS) or Wet Stress (WS) will accumulate once the temperature or soil moisture exceeds DV3 or SM3 respectively. The individual stress indices are combined multiplicatively, indicating that they can operate independently and concurrently. Growth and Stress indices are calculated on a weekly basis and integrated to annual summary variables. This allows the model to capture important seasonal population responses at a temporal scale that is relevant to the dynamics of insect and plant species. An annual value of 100 for the Annual Stress Index is lethal and precludes a species from persisting in a given location. The Weekly and Annual Growth Index (GIW and GIA) represent the potential for population growth and development without considering stresses that may inhibit or completely prohibit population growth. In addition to the temperature and moisture stresses, the potential distribution of *C. capitata* was set to be limited by the minimum length of the growing season measured in degree-days.

The *Compare Locations* model was run using 30-year climate averages from CliMond data centred on 1995 (CM10 World 1995H V2) to fit model parameters and represent the climate suitability for *C. capitata* ^30^. Version 4 of CLIMEX introduces a module that explores climate suitability as a spatio- temporal phenomenon, running the *Compare Years/Locations* model on a time-series of daily values for climate variables. We examined the time series of Annual Growth Index (GIA), stress indices (CS, DS, HS and WS), and the Ecoclimatic Index (EI) values between 1970-2019. The GIA, Temperature Index and Moisture Index values range from 0-100, while stress indices values range between 0- infinity, with values above 100 deemed lethal. EI values range from 0 to 100 and illustrate the overall climatic favourability for the species at specific locations^25^. The Compare Locations function in CLIMEX calculates EI based on 30-years average weather data, while the Compare Years/Locations data calculates EI based on weather data for each individual year in chosen time range. We calculated average EI values for 1970-1979 and 2010-2019 based on EI values returned by Compare Locations/Years function in CLIMEX.

### Parameter fitting

The set of CLIMEX parameters developed by Vera *et al*.^14^ was used as a benchmark for fitting the CLIMEX model. The parameters were carefully revised to ensure they correspond to a plausible biological range of values from experimental work published in the literature. Parameters were fitted by manually, iteratively adjusting the temperature and moisture indices, as well as stress indices, until the locations in marginal *C. capitata* distribution locations corresponded with the geographical distribution simulated by CLIMEX (EI values >= 0.1). We parameterised the model based on CliMond 30-years average climate data centred on 1995^30^. The stress parameters were mostly informed by the distribution data, and the growth parameters were mostly informed by ecophysiological studies and some theoretical considerations (see below).

### Optimal temperature

A study of reproductive and population parameters of *C. capitata* by ^31^ determined that the optimal temperature for development was in the range of 24 - 29 °C. The highest net reproduction rates occurred at 24 °C, while the highest population growth rates were found at 29 °C. A study by Rivnay published in 1950^32^ found that optimal temperatures for egg hatching rates and pupae survival of *C. capitata* was in the range of 25 - 27 °C. These results are consistent with experimental data published by Shoukry and Hafez in 1979^33^, reporting the optimal development temperature of laboratory colonies of *C. capitata* between 25 - 27 °C, when the highest rates of emerging offspring was observed. This value is very close to the optimal temperature of 25 °C proposed in Duyck and Quilici^34^, which was found most favourable for *C. capitata* development time and survival.

### Maximum temperature

Shoukry and Hafez^33^ reported 35 °C as the maximum temperature (Tmax) for *C. capitata* development, similarly to Ricalde *et al*.^10^ who reported 35 °C as the maximum temperature (Tmax) for Brazilian wild populations of *C. capitata*, above which the development of immature stages was inhibited. At 32 °C there was a 68.6% mortality rate for pupae from laboratory colonies^35^.

Duyck and Quilici^34^ also determined that the maximum temperature for *C. capitata* (population from the island of Reunion) was 35 °C; at this temperature, larval stages can barely survive, experiencing 95% mortality.

### Minimum temperature

Accurately estimating the minimum temperature for a species to develop is notoriously difficult. Reports of base temperature in the literature vary depending on several factors, including study location and stage of the fruit fly development. Moreover, recent studies on thermal tolerance and critical minimum and maximum thermal limits^36–38^ have shown that there can be high levels of adaptation and thermal acclimation in local medfly populations, affecting the survival of flies in cold conditions. In Israel the threshold for larval development for *C. capitata*, i.e. the temperature at which they cease to grow, was estimated at between 10-11 °C^32^ and at 8.1 °C in Brazil^10^. We set the minimum temperature for *C. capitata* development (DV0) as 10 °C.

#### Cold stress (CS)

Cold stress studies in literature examined immature stages of *C. capitata* reared in laboratory on artificial diets. The degree-day cold stress (DTCS) was adopted from a previous study^14^ and left at 5 °C with the degree-day cold stress accumulation rate (DHCS) of -0.0015 week^-1^. The cold stress average temperature threshold in our model (TTCSA) is set to 2.5 °C. When temperatures fall below this threshold, stress accumulates at a rate (THSCA) of -0.01 week^-1^.

#### Heat stress (HS)

We set the heat stress value at 37 °C. Literature reports a range of values above 37 °C to induce mortality in medflies^32,39^.

#### Moisture

We left the lower moisture threshold (SM0) set to 0.1^14^ as it reflects the permanent wilting point, typically ca. 10% of soil moisture. The lower and upper optimal moisture parameters (SM1 and SM2) remained 0.3 and 1, respectively (1 indicating 100% soil moisture accumulation), and the limiting high moisture (SM3) parameter was explored to match the distribution of *C. capitata* occurrence location and was set to 1.5, the lower limit for Wet Stress.

#### Dry stress (DS)

Previous study suggests that long periods of water shortage in soils with low water retention capacity in combination with low temperatures are the likely cause of high pupae mortality^40^. Therefore, we set the dry stress threshold moisture level (SMDS) to 0.1 and dry stress accumulation rate (HDS) to - 0.0001 week^-1^.

#### Wet stress (WS)

Some studies have shown a very adverse effect of excessive soil moisture on the development and survival of *C. capitata*, especially at immature stages of development due to potential fruit decomposition (at larval stage) or the pupae’s inability to breathe in highly water-saturated soils. ^32^ reported high mortality rates of the immature *C. capitata* stages associated with excessive soil moisture. More specifically, they found that heavy rainfall, especially when combined with low temperatures, causes high pupae mortality. Flooding of soil caused by heavy rainfall during the rainy season was documented to cause hypoxia in the immature developmental stages of *C. capitata* leading to death^40^ and the precipitation significantly reduces the success of adult emergence^41^.

The wet stress parameter (SMWS) was set to 2 indicating heavy runoff, 100% above soil holding capacity, and the stress accumulation rate (HWS) was set to 0.01 week^−1^.

#### Ecoclimatic Index (EI) calculation

The Ecoclimatic Index (EI) depicts the favourability of the climatic conditions in a given location for the species to grow represented by the Annual Growth Index (GIA) derived from the sum of Weekly Growth Indices (GIW) and representing potential for species to grow (Equation 1). Detrimental effect on species growth during unfavourable seasons is incorporated and summarised in EI as an Annual Stress Index (SI, Equation 2). The EI, a product of GIA and SI (Equation 3), is theoretically scaled between 0 and 100, with an EI of 0 indicating that the location is not favourable for the species’ long-term survival. Very low values of EI indicate marginal climatic suitability. In practice, EI values of 100 are achievable only under stable conditions typically found in equatorial locations.

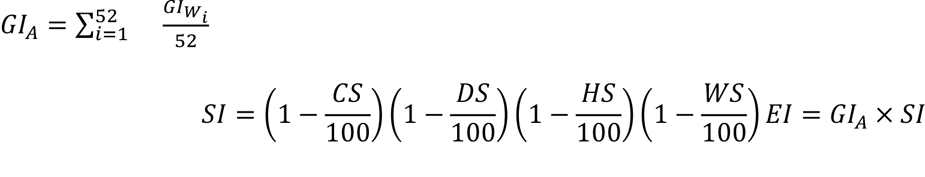

The complete life cycle of *C. capitata* lasts from 20 to 90 days, depending on the prevailing climatic conditions^42^. We set the value of PDD possibly to match an experimental number of generations recorded in locations where we have data. The thermal constant (K) for the total developmental time of *C. capitata* from the populations of Pelotas, Petrolina and Campinas was estimated at 350, 341 and 328 degree days, respectively^10^. We set the required heat sum to complete a generation at 350 degree days.

#### Time-series of climate suitability for *C. capitata*

The CLIMEX Compare Locations/Years module was run on a monthly time series climate database for 1970–2019 to simulate and visualise the seasonal and inter-annual spatio-temporal patterns of climate suitability for *C. capitata* globally. We fitted a linear model using lm() function in R ^17^ at each spatial grid point to the annual CLIMEX index values to estimate the average annual change in EI, GIA, TI, CS, HS and MI across the 50 years in the time series of model outputs. The rate of change over the 50-year timeseries of the model outputs was defined by the slope parameter in the fitted linear model (*a* parameter) and the statistical significance of the slope parameter (p-values) were returned for each grid location and plotted. We assumed that p values < 0.05 indicate significant results.

#### Parameter sensitivity and model uncertainty

CLIMEX V4 ^25^ includes automatic procedures to estimate parametric sensitivity and overall model uncertainty. The aim of this analysis is to draw attention to those parameters whose accurate estimation is important for the model results. These parameters can then be more carefully scrutinised in terms of the reliability with which they have been fitted. Parameters that are sensitive and reliably fitted are of little concern. Conversely, those that are sensitive and uncertainly specified need to be highlighted along with the state variables that are sensitive to these parameters. Some combinations of parameters and state variables are inherently sensitive, and some are inherently insensitive. For example, we expect that the Dry stress variable would be sensitive to the Dry stress threshold (SMDS) and Dry stress accumulation rate (HDS) parameters, but the temperature related variables would be completely insensitive to the dry stress parameters. These expected positive and negative relationships act as a form of validation check to ensure that nothing untoward is occurring in the analysis.

The parametric sensitivity analysis adjusts each model parameter upwards or downwards, and calculates the effect on each of the model variables. Because of anisotropy amongst the model driving variables, the perturbations are fixed using values that reflect the scale of each variable. The calculated parameter sensitivity statistics apply to the climate stations included in the specific simulation. In this case, they reflect the global pattern of sensitivity. In addition to the standard state variables, this analysis includes a variable for the number of climate stations (cells) that have a positive EI value. The calculated sensitivity for this Range Change variable is the proportion of cells that change in suitability for establishment between the two parameter scenarios. We ran the sensitivity analysis for the World using CM10 World 1995H V2 with the rainfed scenario. The results are available in Supporting Information (Table S1).

Where the sensitivity analysis is focused on the parameters, the uncertainty analysis is focused on the results. The uncertainty analysis uses a latin hypercube method to efficiently sample the parametric uncertainty space. The results are an “agreement map”, detailing the proportion of modelled parameter samples that resulted in each cell being classified as being suitable for establishment. Cells that have a high proportion of models indicating potential for survival may be considered to be more robustly modelled as being suitable. In this paper, we ran the uncertainty analysis for the World using CM10 World 1995H V2 with the rainfed scenario.

## Discussion

Our analysis highlights the ongoing effects of climate change which is steadily shifting the potential geographic distribution of *C. capitata* towards regions considered previously suitable for ephemeral populations only. The statistically significant increasing climate suitability for *C. capitata* in Europe and North America from the 1970’s to the 2010’s is marked (Figure 4). This polewards expansion parallels a substantial increase in altitudinal range in tropical Africa and South America (Figure 7). These changes are mostly driven by increasing temperatures, elevating the Temperature Index (TI) and decreasing Cold Stress (CS). The modelled range decreases follow decreasing soil moisture suitability due to decreased rainfall or rising evapotranspiration rates in semi-arid regions, for example in southern Africa and central Australia (Figure 7).

Our analysis reveals that persistence of *C. capitata* in some central and northern parts of Europe derived from the GBIF database, for example in Romania and around Lyon valley, would require biologically radical adaptation to the cold stresses currently limiting *C. capitata* persistence in this area. *C. capitata* may be able to overwinter in sheds, and protected anthropogenic places not exposed to lethal winter temperatures ^20^. Indeed, recent studies conducted in the area of Vienna in Austria revealed that *C. capitata* can overwinter in human made shelters but not in open field conditions.

Conversely, our results reveal that the persistence of *C. capitata* in northern Italy and the Paris Basin is increasingly likely, which is consistent with the simulations of Gilioli et al.^43^. Locations with documented multi-season populations in Germany and Austria also observe a positive trend in TI, indicating the conditions are becoming increasingly favourable. With the ongoing positive trend in climatic suitability changes in this area, *C. capitata* may yet be able to overwinter in these locations under field conditions^44^.

In California the main limitation to *C. capitata* development is a lack of sufficient rainfall and consequently, soil moisture conditions. Our results represent a composite scenario, where the majority of the state appears to be equipped for irrigation^23^ and a sufficient top-up irrigation is available throughout the season, removing the dry stress that would otherwise limit *C. capitata* survival. Our analysis shows that most of California has witnessed a consistent improvement in conditions suitable for *C. capitata*. Additionally, over the past fifty years, the entire northern limit of the species’ suitability expanded poleward, while some areas in the south became unsuitable due to heat stress (HS).

This expansion of the climatic niche into higher latitudes and altitudes could have significant implications for the horticultural industries. Firstly, the previously unaffected regions may now become vulnerable to pest infestations. Secondly, the length of the suitable conditions within a season may increase, which could result in more generations being completed in one season. Increased population densities later in the season may lead to significant infestation rates on later ripening hosts. Thirdly, where the climate suitability changes from ephemeral (relying on source-sink metapopulation dynamics to re-invade) to suitable for year-round persistence, the starting population of flies at the beginning of spring could increasedramatically.

Our comparisons between the 1970s and the 2010’s highlights that climate is changing at a rate that is statistically significant in terms of geographical pest risks. The historically popular climate datasets centred on the 1970’s are now outdated. Hence, there is a pressing need to use recent climatologies in species niche modelling in order to correctly reflect the species climatic range. Climatologies typically span 30 years, so that inter-decadal variation is ameliorated. Given the observed rate of change in climate and associated pests risks, a 20-year climatology may be a better suited standard to use to model contemporary risks. Pest risk analysts and managers would also be well advised to consider future climate scenarios to inform their emerging risks.

Our results demonstrating statistically significant trends in climate suitability for *C. capitata* in Europe and California echo those for *Bemisia tabaci* in Eastern Africa^3^, adding to the body of evidence linking observed climate changes to changes in pest dynamics. These findings underscore the urgency of addressing this factor in preparedness for potential future pest invasions. Our research emphasizes the importance of basing biosecurity decisions on modelling that uses up-to-date climatic information and takes into account that pest risk areas are shifting. With ongoing shifts in climate suitability, it becomes crucial to understand the rate at which pest suitability is changing and the timeframe in which previously unsuitableareas may become prone to successful pest establishment. This knowledge is essential for effective pest management strategies as we navigate the evolving challenges of a changing climate.

## Acknowledgements

The authors gratefully acknowledge funding from the European Union’s Horizon 2020 research and innovation programme under grant agreement No 818184. We extend our sincere thanks to Adam and Josep Jaques Miret and Andrea Sciarretta for their expert insight into the assessment of distribution data in Spain and Italy.

## Data availability statement

The occurrence data used in this study are openly available on Figshare at doi:10.6084/m9.figshare.23721477.

