## Supplementary Material for "Evidence that recent climatic changes have expanded the potential geographical range of the Mediterranean fruit fly"

**Szyniszewska et al.**

### Supporting information

| Parameter description | Parameter | Low | Actual | High | Run | CD Change | CS Change | DS Change | EI Change | GI Change | HS Change | MI Change | Range Change | TI Change |
|---|---|---|---|---|---|---|---|---|---|---|---|---|---|---|
| Dry Stress Threshold | SMDS | 0 | 0.1 | 0.2 | 15 | 20.32 | 0 | 68.52 | 11.94 | 0 | 0 | 0 | 15.75 | 0 |
| Dry Stress Rate | HDS | -0.06 | -0.05 | -0.04 | 16 | 0.26 | 0 | 9.59 | 1.84 | 0 | 0 | 0 | 2.94 | 0 |
| Limiting low moisture | SM0 | 0 | 0.1 | 0.2 | 1 | 0 | 0 | 0 | 4.99 | 6.9 | 0 | 15.3 | 1.27 | 0 |
| Cold Stress Degree-day Threshold | DTCS | 4 | 5 | 6 | 9 | 0.7 | 8.1 | 0 | 1.27 | 0 | 0 | 0 | 1.05 | 0 |
| Wet Stress Threshold | SMWS | 1.4 | 1.5 | 1.6 | 17 | 2.34 | 0 | 0 | 2.76 | 0 | 0 | 0 | 0.81 | 0 |
| Cold Stress Degree-day Rate | DHCS | -0.0018 | -0.0015 | -0.0012 | 10 | 0.08 | 4.9 | 0 | 0.84 | 0 | 0 | 0 | 0.72 | 0 |

| | | | | | | | | | | | | | | |
|---|---|---|---|---|---|---|---|---|---|---|---|---|---|---|
| Lower optimal moisture | SM1 | 0.2 | 0.3 | 0.4 | 2 | 0 | 0 | 0 | 4.37 | 5.4 | 0 | 12.47 | 0.32 | 0 |
| Wet Stress Rate | HWS | 0.012 | 0.015 | 0.018 | 18 | 0.06 | 0 | 0 | 0.91 | 0 | 0 | 0 | 0.3 | 0 |
| Limiting high moisture | SM3 | 1.4 | 1.5 | 1.6 | 4 | 0 | 0 | 0 | 3.72 | 4.2 | 0 | 7.52 | 0.2 | 0 |
| Heat Stress Temperature Threshold | TTHS | 38 | 39 | 40 | 13 | 0 | 0 | 0 | 0.43 | 0 | 15.36 | 0 | 0.18 | 0 |
| Limiting low temperature | DV0 | 9 | 10 | 11 | 5 | 0 | 0 | 0 | 1.73 | 2.3 | 0 | 0 | 0.16 | 3.6 |
| Cold Stress Temperature Threshold (Average) | TTCSA | 1.5 | 2.5 | 3.5 | 11 | 0 | 1.6 | 0 | 0.32 | 0 | 0 | 0 | 0.16 | 0 |
| Limiting high temperature | DV3 | 36 | 37 | 38 | 8 | 0 | 0 | 0 | 0.68 | 1.3 | 0 | 0 | 0.1 | 3.7 |
| Lower optimal temperature | DV1 | 27 | 28 | 29 | 6 | 0 | 0 | 0 | 2.05 | 2.5 | 0 | 0 | 0.08 | 5.2 |
| Upper optimal temperature | DV2 | 32 | 33 | 34 | 7 | 0 | 0 | 0 | 1.49 | 2.2 | 0 | 0 | 0.08 | 4.6 |
| Upper optimal moisture | SM2 | 0.9 | 1 | 1.1 | 3 | 0 | 0 | 0 | 2.65 | 3 | 0 | 8.24 | 0.04 | 0 |
| Heat Stress Temperature Rate | THHS | 0.008 | 0.01 | 0.012 | 14 | 0 | 0 | 0 | 0.07 | 0 | 3.31 | 0 | 0.02 | 0 |
| Degree-days per Generation | PDD | 280 | 350 | 420 | 21 | 0 | 0 | 0 | 0.08 | 0 | 0 | 0 | 0.02 | 0 |

| | | | | | | | | | | | | | | |
|---|---|---|---|---|---|---|---|---|---|---|---|---|---|---|
| Cold Stress Temperature Rate (Average) | THCSA | -0.012 | -0.01 | -0.008 | 12 | 0 | 0.1 | 0 | 0 | 0 | 0 | 0 | 0 | 0 |
| Hot-Wet Temperature Threshold | TTHW | 0 | 0 | 1 | 19 | 0 | 0 | 0 | 0 | 0 | 0 | 0 | 0 | 0 |
| Hot-Wet Moisture Threshold | MTHW | 0 | 0 | 0.1 | 20 | 0 | 0 | 0 | 0 | 0 | 0 | 0 | 0 | 0 |

Table S1. The CLIMEX sensitivity analysis outputs, run with parameters above or below the original parameter values.

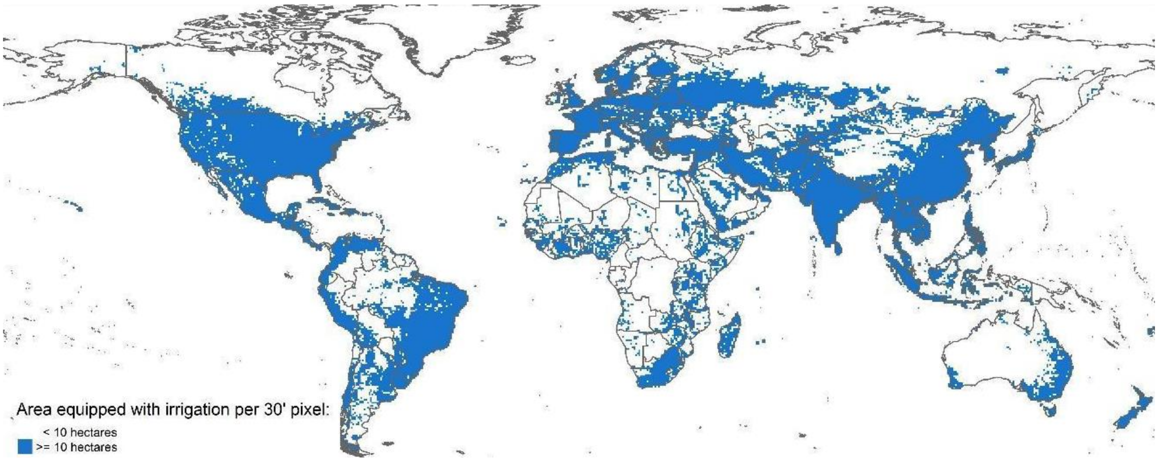


Figure S1. Irrigation map created to create CLIMEX composite model layers accounting for scenarios with or without irrigation. The classification was based on the Global Map of Irrigation Areas version 5.0 (FAO) using the area equipped for irrigation expressed in hectares layer. The layer was resampled from 10-minute to 30-minute spatial resolution to match the CliMond climate dataset. Areas with less than 10 hectares of land equipped in irrigation were classified as non-irrigated (white), and those with the irrigation at or above 10 hectares per 30-minute pixel were classified as irrigated (blue).

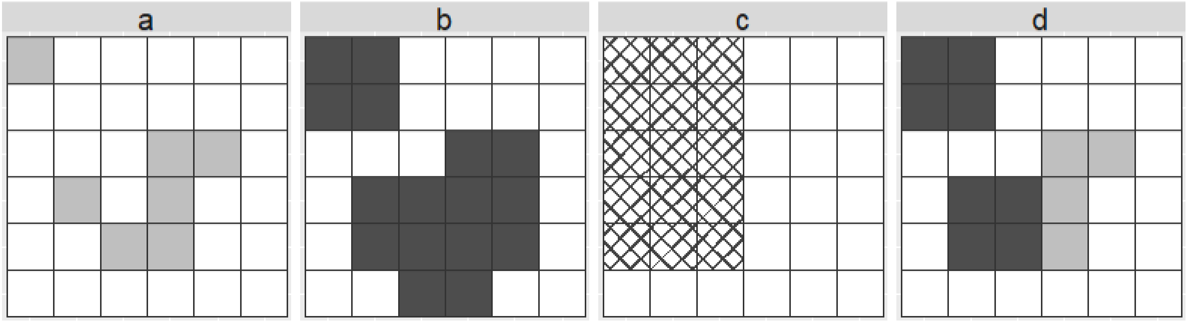


Figure S2. Process building composite maps (adopted from Yonow et al.). a. Under natural rainfall scenarios shaded with CLIMEX index values over 0; b. Areas suitable under an irrigation scenario; c. Irrigated areas; d. Composite map of the areas of ‘a’ and ‘b’, where Climex values from ‘a’ are taken in locations not equipped in irrigation, and CLIMEX values from ‘b’ are taken in areas equipped in irrigation.

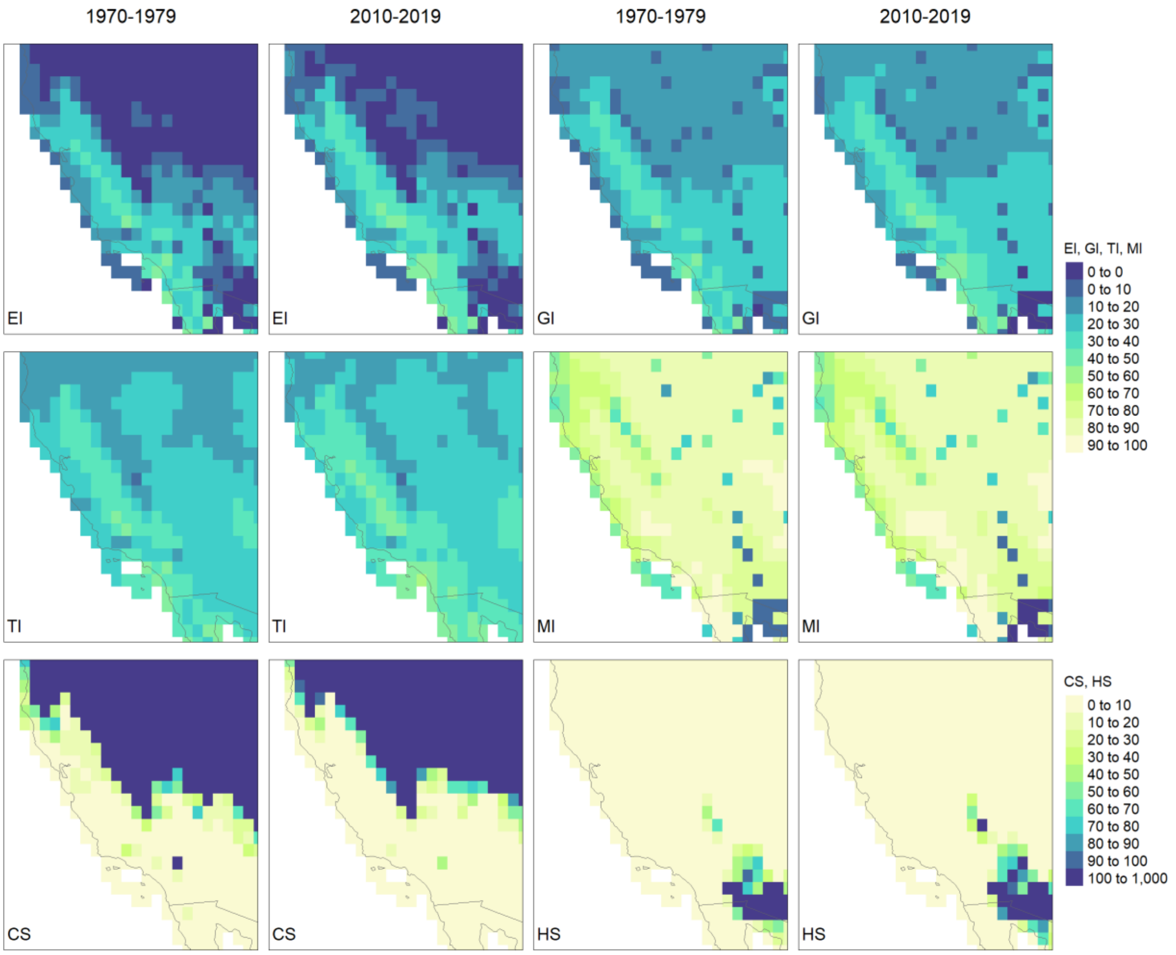


Figure S3. CLIMEX model results representing 10-year average values of Ecoclimatic index (EI), Growth Index (GI), Temperature Index (TI), Moisture Index (MI), Cold Stress (CS) and Heat Stress (HS) in California for 1970-1979 (1st and 3rd column) and 2010-2019 (2nd and 4th column). The Compare Locations/Years function was run utilising CRU climate data returning annual index and stress values.

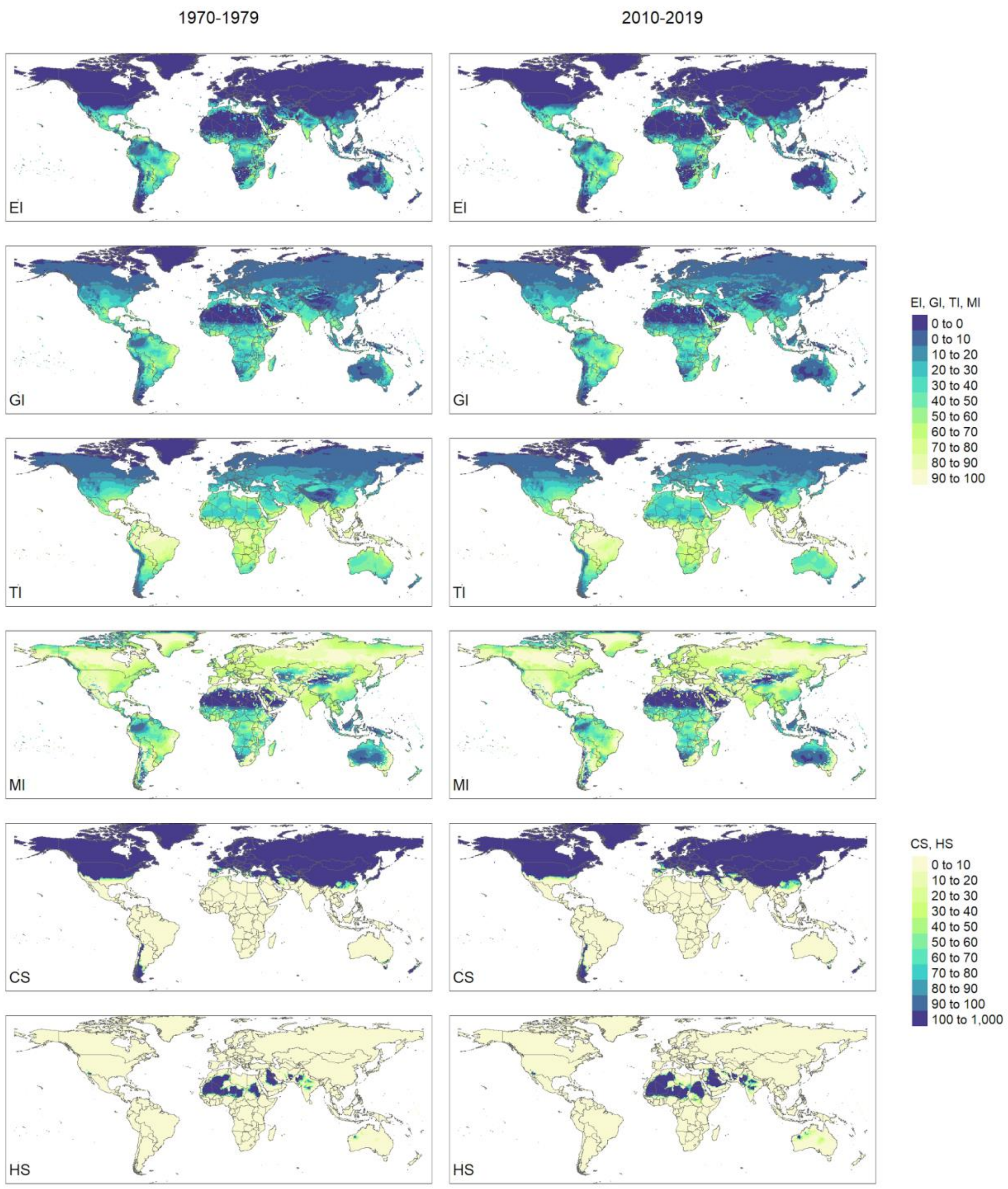


Figure S4. CLIMEX model results representing 10-year average values of Ecoclimatic index (EI), Growth Index (GI), Temperature Index (TI), Moisture Index (MI), Cold Stress (CS) and Heat Stress (HS) globally for 1970-1979 (left column) and 2010-2019 (right column). The Compare Locations/Years function was run utilising CRU climate data returning annual index and stress values.

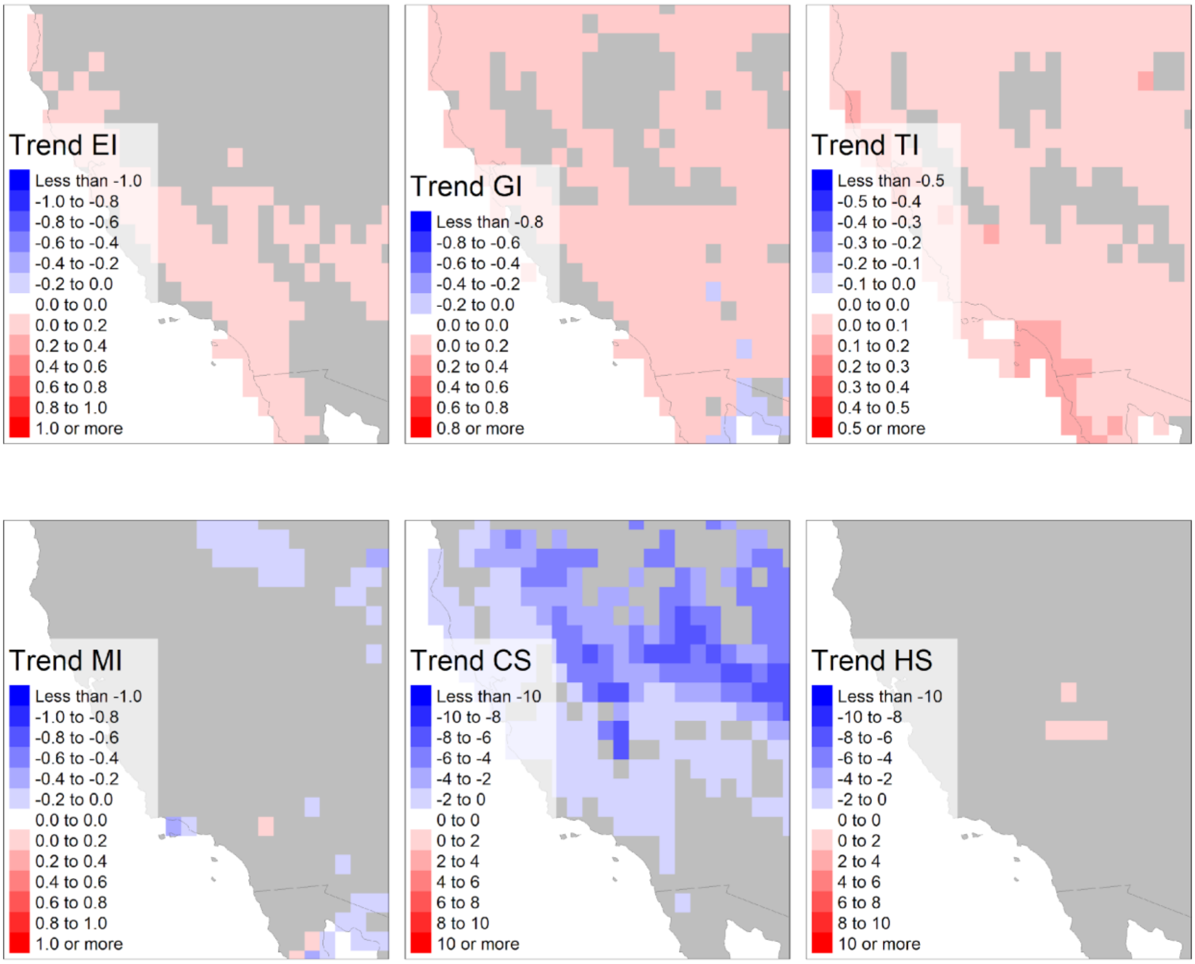


Figure S5. Annual rate of change (trend) values for medfly in California returned by lm() function in R statistical programming software run on annual values returned by Compare Locations/Years function for 1970-2019 for six CLIMEX indices: Ecoclimatic index (EI), Growth Index (GI), Temperature Index (TI), Moisture Index (MI), Cold Stress (CS) and Heat Stress (HS).

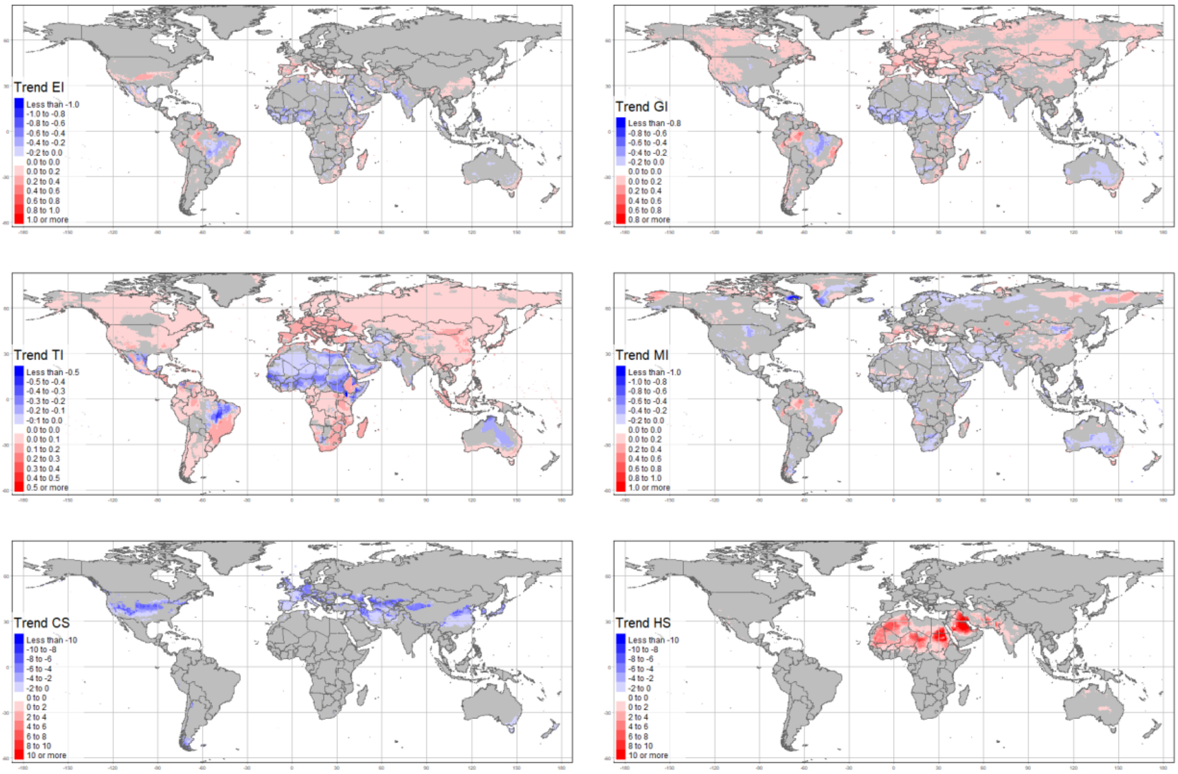


Figure S6. Global representation of the significance ($p < 0.05$) of the slope parameter value in a linear model fitted to the 50-year time-series (1970-2019) of the Ecoclimatic Index (EI), Growth Index (GI), Temperature Index (TI), Moisture Index (MI), Cold Stress (CS) and Heat Stress (HS) CLIMEX model outputs.

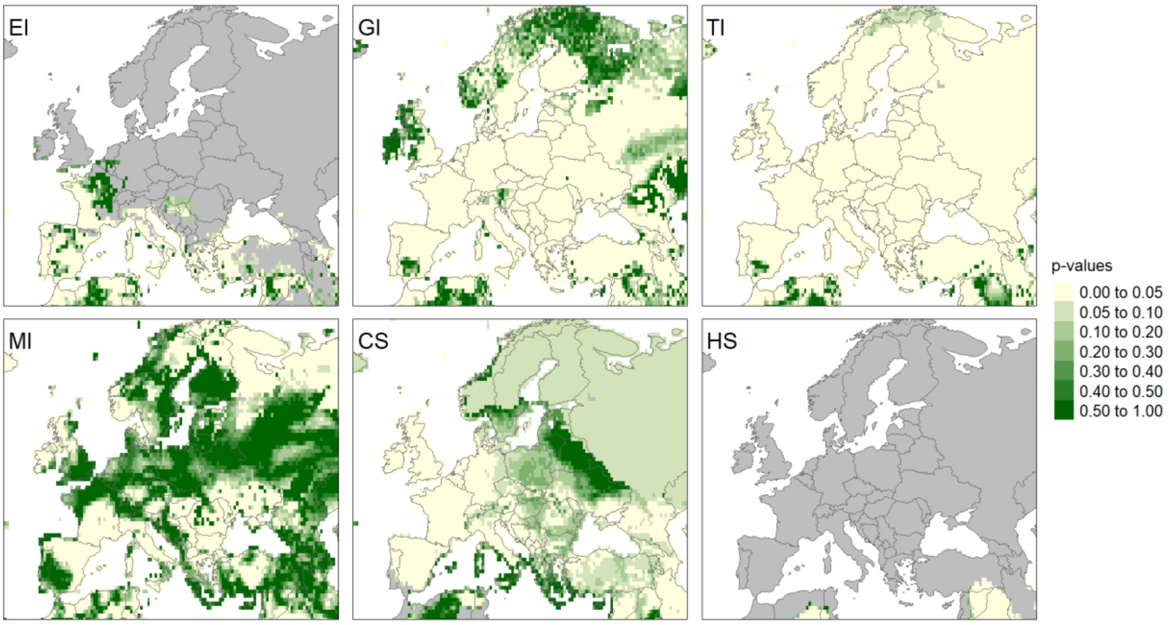


Figure S7. Statistical significance represented by p-values of calculated 50-year linear trends for the 50-year time series of six CLIMEX index values: Ecoclimatic index (EI), Growth Index (GI), Temperature Index (TI), Moisture Index (MI), Cold Stress (CS) and Heat Stress (HS) in Europe.

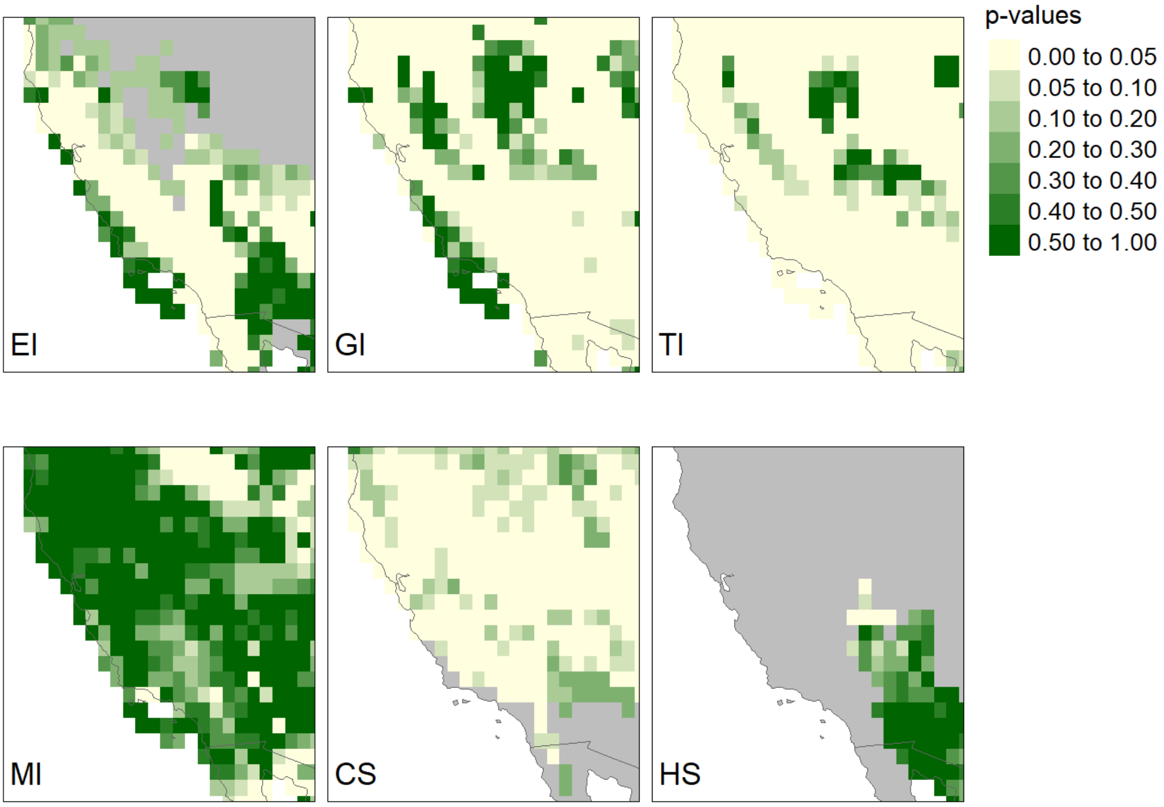


Figure S8. The significance (*p*) value of the slope parameter value in a linear model fitted to the 50-year time-series (1970-2019) of the Ecoclimatic Index (EI), Growth Index (GI), Temperature Index (TI), Moisture Index (MI), Cold Stress (CS) and Heat Stress (HS) CLIMEX model outputs. Results represent California, USA and surrounding areas.

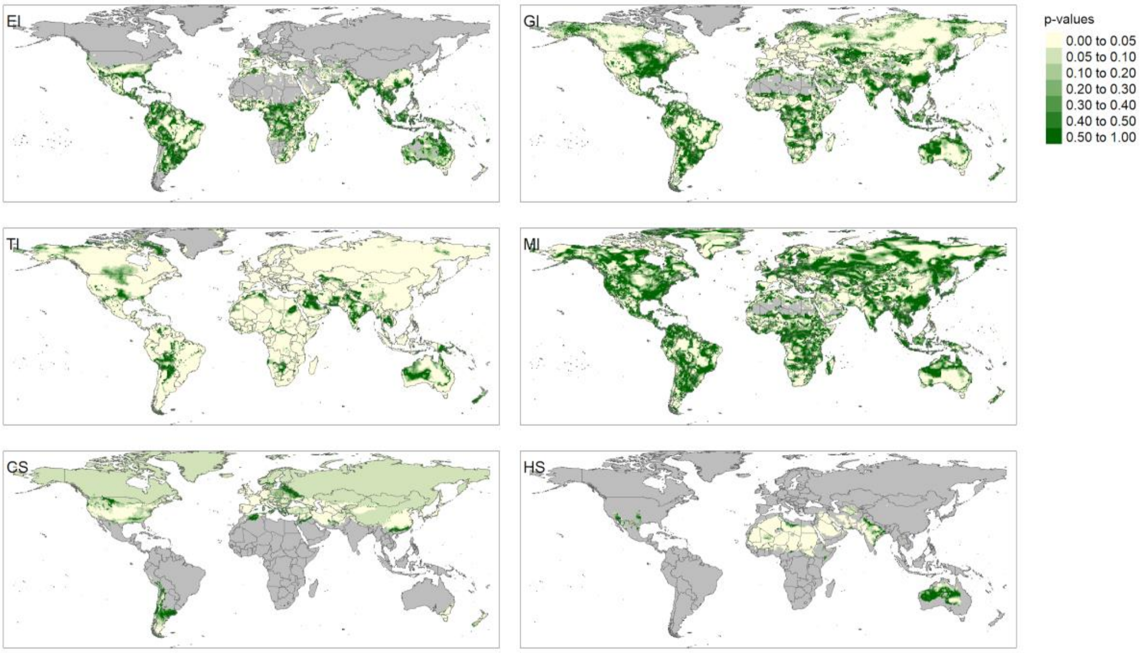


Figure S9. Global representation of the significance (p) value of the slope parameter in a linear model fitted to the 50-year time-series (1970-2019) of the Ecoclimatic Index (EI), Growth Index (GI), Temperature Index (TI), Moisture Index (MI), Cold Stress (CS) and Heat Stress (HS) CLIMEX model outputs.

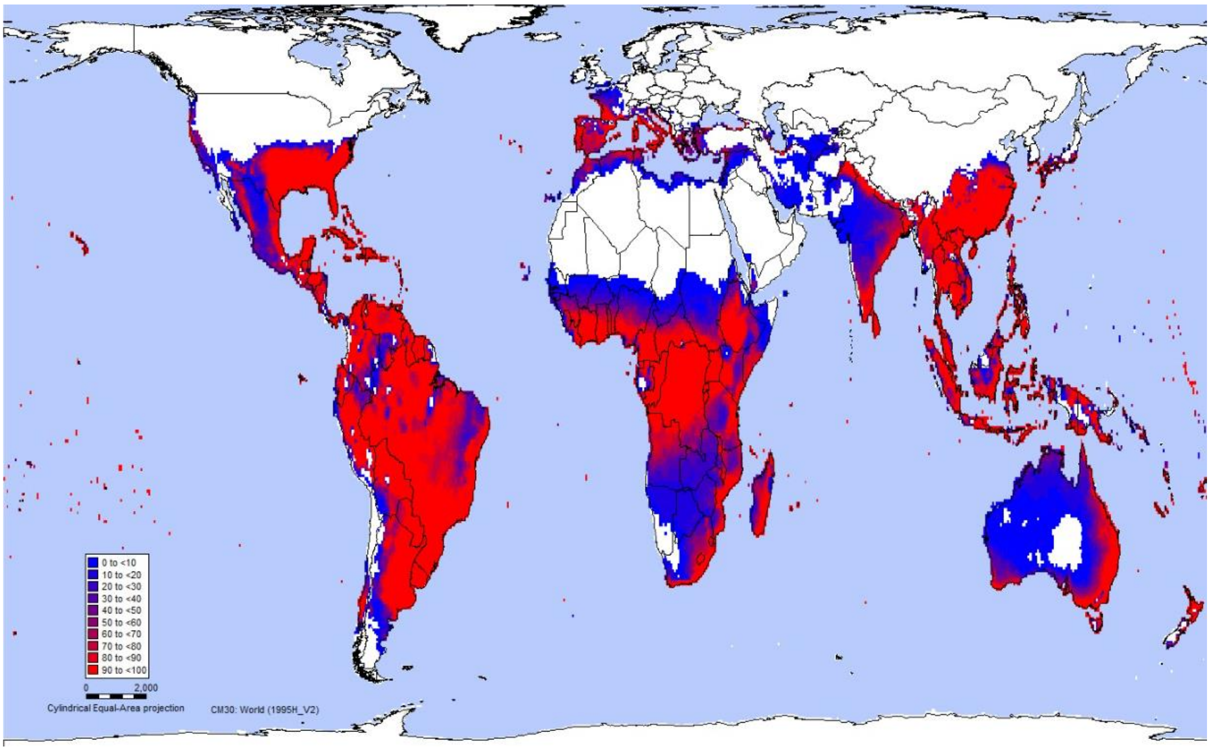


Figure S10. CLIMEX model prediction uncertainty map.
